# The effects of pks^+^ *Escherichia coli* and bile acid in colorectal tumorigenesis among people with cholelithiasis or cholecystectomy

**DOI:** 10.1101/2022.12.07.519553

**Authors:** Si-Yuan Pan, Cheng-Bei Zhou, Jia-Wen Deng, Yi-Lu Zhou, Zhu-Hui Liu, Jing-Yuan Fang

## Abstract

Patients with cholelithiasis (CL) or cholecystectomy (CE) would have more chances to get colorectal adenoma (CRA) or cancer (CRC). To figure out the effect of gut microbiota and bile acid on colorectal neoplasm in CL and CE patients, we executed a retrospective observational study recruited 463 volunteers, including 182 people with normal gallbladder (Normal), 135 CL and 146 CE patients. The discovery cohort was established to explore the difference of gut microbiota through 16S rRNA sequencing. The validation cohort aimed to verify the results of sequencing through qPCR. Through this research, significant enrichment of *Escherichia coli* was found in patients with cholelithiasis or cholecystectomy both in discovery cohort (*P*_Normal-CL_=0.013; *P*_Normal-CE_=0.042) and in validation cohort (*P*_Normal-CL_<0.0001; *P*_Normal-CE_<0.0001). The relative abundance of *Escherichia coli* was also increased in CRA and CRC patients (in discovery cohort, *P*_HC-CRA_=0.045, *P*_HC-CRC_=0.0016; in validation cohort, *P*_HC-CRA_=0.0063, *P*_HC-CRC_=0.0007). Pks^+^*Escherichia coli* was found enriched in CL and CE patients in validation cohort (*P*_Normal-CL_<0.0001; *P*_Normal-CE_<0.0001). Through KEGG analysis in discovery cohort, the differences of bile acid metabolism were revealed (Ko00120 primary bile acid biosynthesis *P*=0.014; Ko00121 secondary bile acid biosynthesis *P*=0.010). In validation cohort, we also found the elevation of serum total bile acid of CE patients (*P*<0.0001). And the level of serum total bile acid was found associated with the relative abundance of pks^+^ *Escherichia coli* (r=0.1895, *P*=0.0012). In one word, our research found that *Escherichia coli*, especially pks^+^ species, was enriched in CL and CE patients. Pks^+^ *Escherichia coli* and bile acid metabolism were associated with CRA and CRC in people after cholecystectomy.

## Introduction

Changes of dietary habits in recent years lead to rapid growth of gallbladder disorders worldwide(1). Cholecystectomy has been an accepted management for gallbladder disorders especially for severe cholelithiasis(2, 3). However, the long-term pathophysiological changes among patients after cholecystectomy remain unclear. Studies have found that compared to those with normal gallbladders, the patients with cholecystectomy would be prone to developing colorectal cancer (CRC) or colorectal adenoma (CRA)(4–6). Meanwhile, it has been reported that patients with gallstone but without cholecystectomy also had a higher incidence of CRC and CRA(4, 7, 8).

Previous studies have figured out that the composition and excretion rhythms of bile acids were different among people with different gallbladder conditions(9–12). The changes of bile acid metabolism in intestinal tract, especially the enrichment of secondary bile acids, were proved to stimulate the malignant transformation of intestinal epithelial cells and lead to the development of CRC(11). It was speculated that the disorders of bile acids could contribute to the occurrence and development of CRA and CRC in patients with cholelithiasis or cholecystectomy.

There have been intensive studies on relationship between gut microbiota and CRC(13, 14). The roles of some bacteria, such as *Fusobacterium nucleatum*(15), *Escherichia coli*(16, 17) and *Bacteroides fragilis*(18), had been pointed out. However, few studies have focused on the effects of the gut microbiota on bile acid metabolism or gallstone formation, and whether they would eventually cause further consequences including the increased incidence of colorectal tumors.

The purpose of current study is to identify the specific bacteria which contribute to the higher risk of CRA and CRC among patients with cholelithiasis and cholecystectomies, and to figure out the effect of gut microbiota and bile acid on the development of colorectal neoplasm in such cases.

## Methods

### Study design and cohort participants

We conducted a retrospective study among patients with different gallbladder conditions in Renji Hospital, Shanghai Jiao-Tong University School of Medicine. A total of 463 volunteers were recruited, including 182 patients with normal gallbladder (Normal), 135 patients with cholelithiasis (CL) and 146 patients with cholecystectomy (CE), and were divided into two cohorts. To further explore the differences of tumor characteristics and bile acid metabolism among Normals, CLs and CEs, we also searched the sample database of CRA and CRC patients from 2018 to 2022 in Renji Hospital and 1,481 patients were involved (675 Normals, 526 CLs and 280 CEs).

The protocol was approved by the Ethics Committee of Renji Hospital (#2022-065-B). The study flowchart was shown in Figure 1. Inclusion and exclusion criteria were described in detail in Table 1.

**Figure 1.**
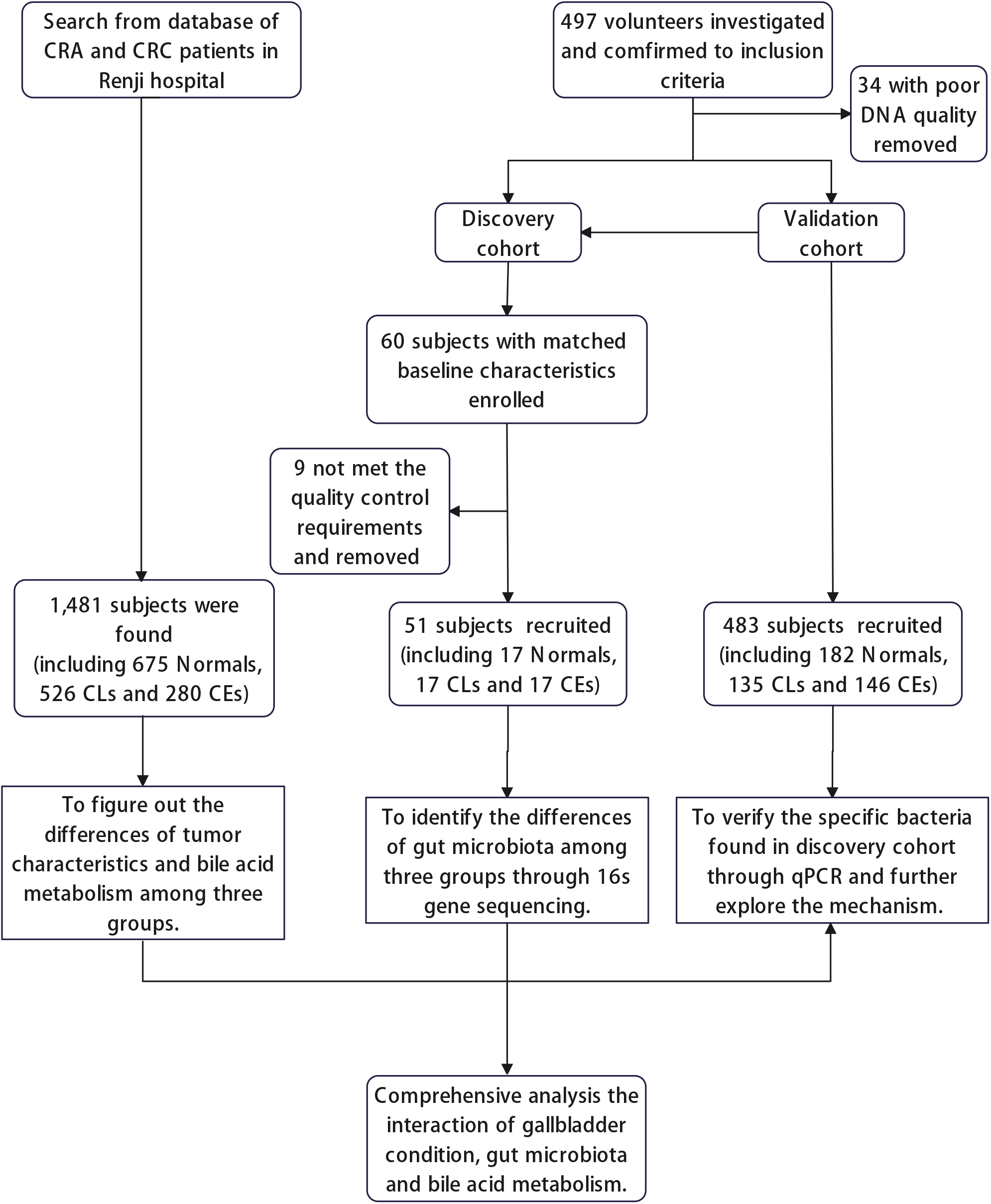
Study work flowchart.

**TABLE 1.** The inclusion and exclusion criteria of our retrospective study

| Inclusion criteria | Exclusion criteria |
| --- | --- |
| (1) Aged between 40 and 85 years. | (1) History of CRC, Lynch syndrome, Peutz–Jeghers syndromes, familial adenomatous polyposis, inflammatory bowel disease, or intestinal tuberculosis. |
| (2) Patients with a clear history of cholecystectomy or had undergone gallbladder imaging (including abdominal ultrasound, computed tomography scan or magnetic resonance imaging) within 1 year prior to colonoscopy or surgery and the result of the gallbladder examine was conformed to be Normal (Normal), cholelithiasis (CL) or after cholecystectomy (CE). | (2) History of gastrointestinal surgery. |
| (3) The results of colonoscopy, surgery or pathology were confirmed to be normal (HC), colorectal adenoma (CRA) or colorectal cancer (CRC). | (3) History of diabetes, chronic liver disease and other disease which would affect bile acid metabolism. |
|  | (4) History of other malignant tumors. |
|  | (5) Usage of any of the following medicines 3 month before colonoscopy or surgery: antibiotics, probiotics, choleretic drugs, glucocorticoid, immunosuppressors, statins, chemotherapy drugs, targeted therapy or immunotherapy drugs. |
|  | (6) Unable to complete the questionnaire, provide clinical information and qualified stool specimens. |

### Colonoscope, Surgery, histopathology, Sample Collection and DNA extraction

We recruited individuals with or without symptoms of digestive tract in endoscopy center and general surgery ward of Renji Hospital. After adequate bowel preparation, all colonoscopies were conducted by experts and reached the ileocecal valve with enough observation time (withdrawal time more than 6 min). All colorectal tumors of recruited patients were removed through surgery or endoscopic resection by experienced surgeons or experts, and were immediately sent for pathological examination. All colonoscope experts, surgeons and pathologists involved were blinded to the research details.

In the discovery and validation cohort, at least 1.0g of stool samples were collected in the germ-free containers before bowel preparation for the colonoscope and surgery. All the stool samples were immediately stored in −20°C and transferred to −80°C refrigerator within 48h. QIAamp PowerFeacl DNA Kit (Qiagen, Hilden, Germany) was used to extract stool genomic DNA. The concentration of DNA was measured by a Nanodrop 2000 instrument (Thermo Scientific, Waltham, MA, USA). All extracts were preserved at −20°C before the subsequent experiment.

### Subject allocation

The patients were divided into three groups according to the condition of gallbladder. The patients with normal gallbladder (Normal) were defined to have a normal gallbladder by abdominal ultrasound within 1 year before colonoscope or surgery and had no history of gallbladder diseases. The patients with cholelithiasis (CL) were found to be detected gallbladder stones by ultrasound, CT or MRI within 1 year prior to colonoscope or surgery. CE group was defined as people underwent cholecystectomy more than 1 year before colonoscope or surgery. Through the results of colonoscope, surgery and pathology, the patients were divided into another three groups, including healthy control (HC), colorectal adenoma (CRA) and colorectal adenocarcinoma (CRC). The CRC stage was assessed by the TNM system according to American Joint Committee on Cancer version 8. 2017(19).

For further research and subgroup analysis, we divided those patients through both gallbladder condition and the results of colonoscope or pathology. NHC refers to volunteers both with a normal gallbladder and a normal colonoscope result. NCRA refers to CRA patients with normal gallbladder, and NCRC refers to CRC patients with normal gallbladder. In patients with cholelithiasis (CL), participants were also divided to healthy control (CLHC), colorectal adenoma (CLCRA) and colorectal adenocarcinoma (CLCRC). Patients with cholecystectomy (CE) were also separated into CEHC (CE patients with normal colonoscope result), CECRA (CE patients with CRA) and CECRC (CE patients with CRC).

### 16S rRNA gene sequencing

The V3-V4 variable region of bacterial 16S rRNA was amplified via PCR using primers (338F, 5’-ACTCCTACGGGAGGCAGCAG-3’; 806R,5’-GGACTACHVGGGTWTCTAAT-3’) with TransStart Fastpfu DNA Polymerase. After purified via gel extraction (AxyPrep DNA GelExtraction Kit, Axygen Biosciences, Union City, CA, USA), the amplicons were sequenced on the Miseq platform (Illumia, San Diego, California, USA) in equivalent concentrations.

### Quantitative Polymerase Chain Reaction (qPCR)

We verified the potential differences of gut microbiota, which were found using 16S rRNA Illumina MiSeq sequencing, in validation cohort by quantitative polymerase chain reaction. 10 μL reaction volume of TB Green^®^ Premix Ex TaqTM II (TaKara, Japan), which included 40ng fecal DNA, was used to measure the relative abundance of *Escherichia Coli*, pks^+^ *Escherichia coli* and cdt^+^ *Escherichia coli* according to the reaction conditions: 95°C for 30s (denaturation), a total of 45 cycles of 95°C for 5s (annealing) and 60°C for 30s (extension). The primers were listed in the following list (Supplementary Table 1) and total bacterial DNA was determined by 16s rRNA. The single defined peak of the melt curve certifies the specificity of each prime (Supplementary Figure 1). Subsequently, the relative abundance of targeted gut microbiota was evaluated by 2^-ΔCt^ (ΔCt= the average Ct of targeted microbiota-the average Ct of 16S rRNA).

### Statistical analysis

The analysis of 16S rRNA gene sequencing raw data were based on the Quantitative Insights Into Microbial Ecology platform (QIIME, V.1.9.1). The UPARSE platform (V.7.0.1090, http://drive5.com/uparse/) helped to clustered operational taxonomic units (OTUs) at the similarity of 97%. The identified taxonomy was based on the database Silva (V.138, http://www.arb-silva.de). Pan, core analysis and rank-abundance curve were used to prove that the sample size and sequencing volume were sufficient (Supplementary Figure 2). Ace, Shannon, Sobs and Chao indexes were estimated to assess species diversity and richness (Supplementary Figure 3). LEfSe analysis was used to discover different species (Figure 2c, f). KEGG function prediction analysis was used through Tax4Fun to explore the possible mechanism (Figure 4a–b).

**Figure 2.**
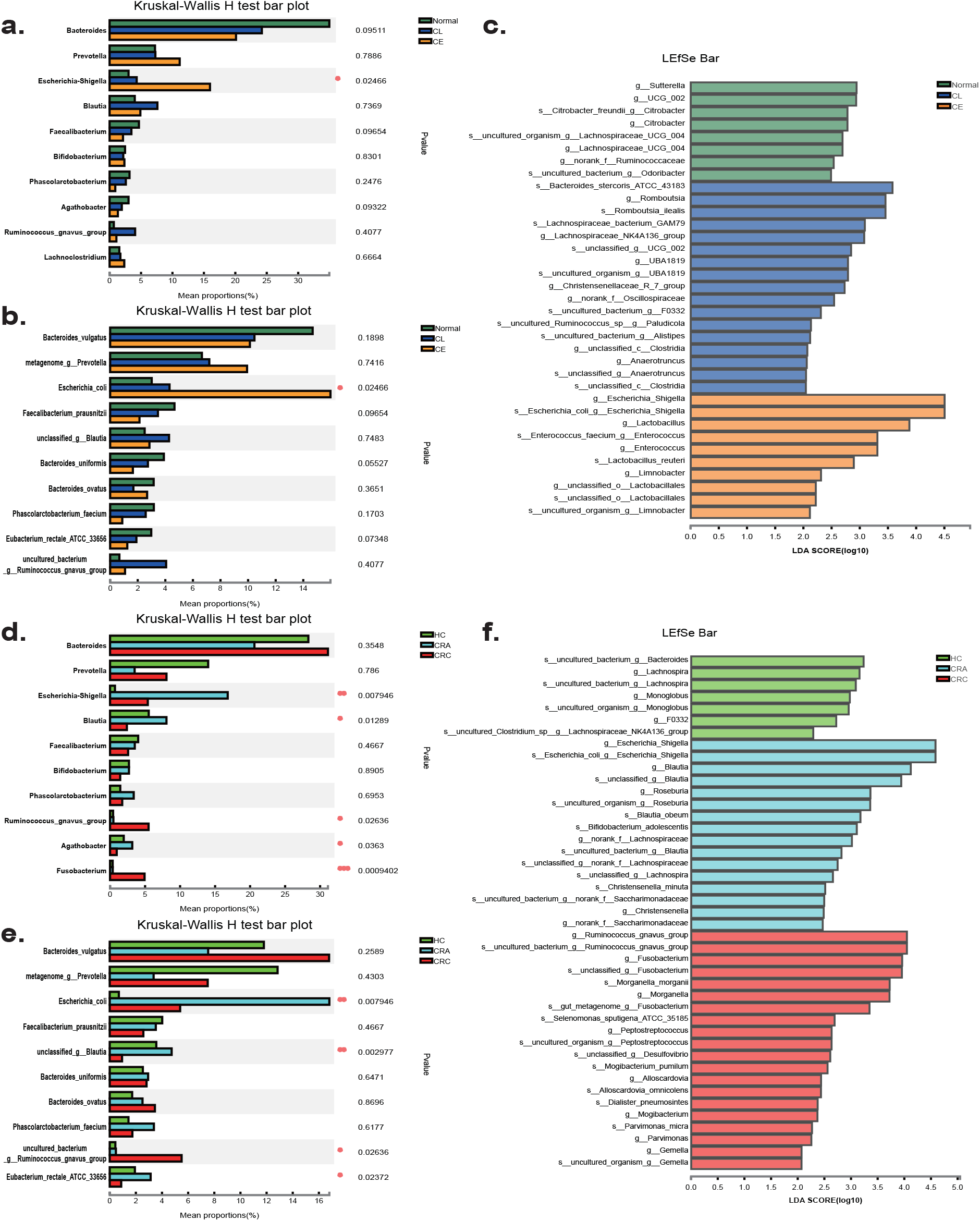
The 16S rRNA Illuminate sequencing result in discovery cohort. (a)-(b) The differences of sequencing proportion in genus(a) and species(b) levels in stool microbiota among Normal, CL and CE patients (Genus: *Escherichia-Shigella*, *P*=0.025; Species: *Escherichia Coli*, *P*=0.025). (c) LEfSe analysis from genus to species among Normal, CL and CE patients in discovery cohort. (d)-(e) The differences of sequencing proportion in genus(d) and species(e) levels in stool microbiota among HC, CRA and CRC patients (Genus: *Escherichia-Shigella*, *P*=0.0079; Species: *Escherichia Coli*, *P*=0.0079). (f) LEfSe analysis from genus to species among HC, CRA and CRC patients in discovery cohort. * *P*<0.05, ** *P*<0.01, *** *P*<0.001

**Figure 3.**
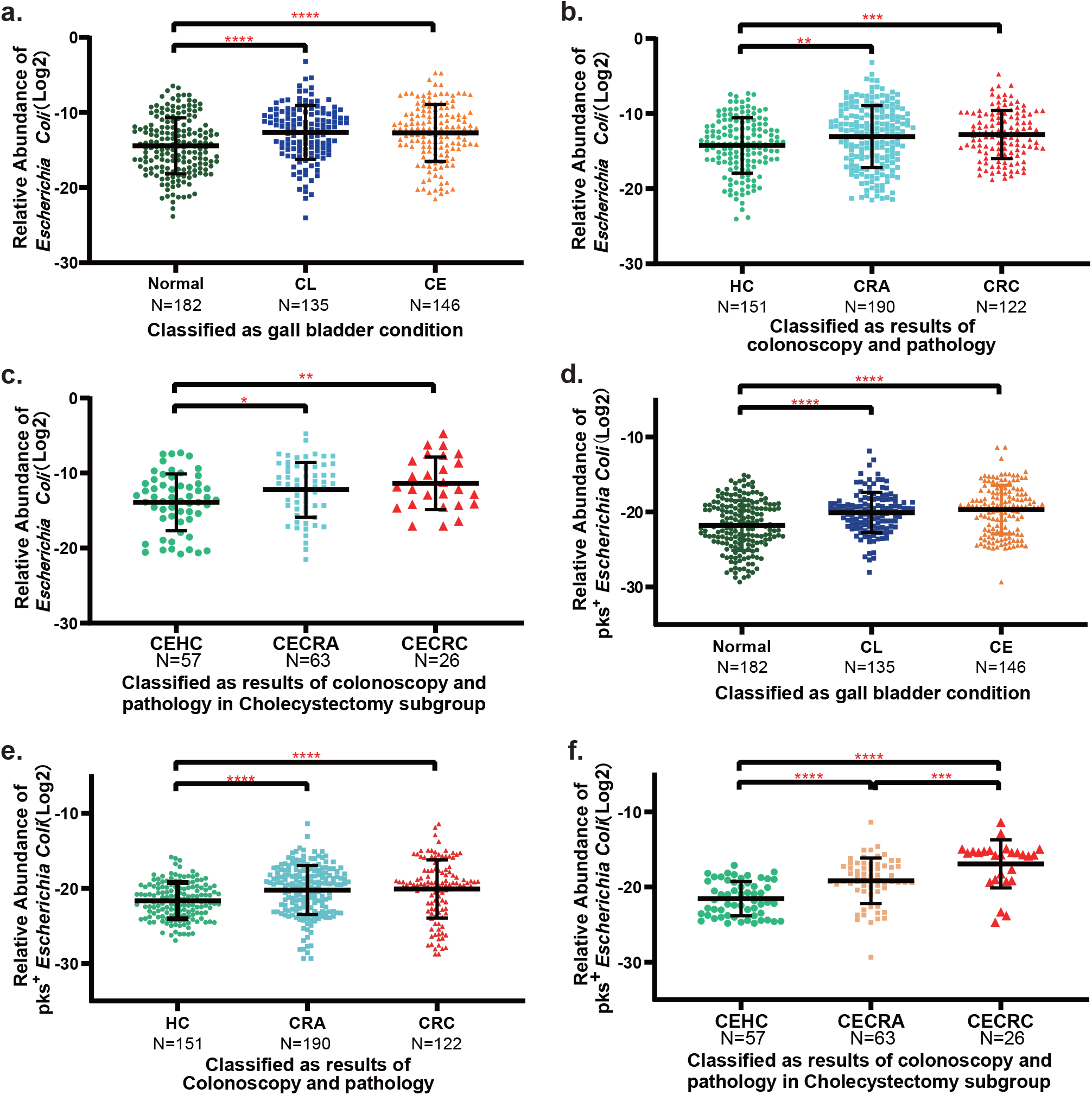
The relative abundance of *Escherichia Coli* and pks^+^ *Escherichia Coli* in validation cohort by qPCR. (a) The difference of the relative abundance of *Escherichia Coli* among Normal, CL and CE patients (*P*_Normal-CL_<0.0001, *P*_Normal-CE_<0.0001). (b) The difference of the relative abundance of *Escherichia Coli* among HC, CRA and CRC patients (*P*_HC-CRA_=0.0063, *P*_HC-CRC_=0.0007). (c) The difference of the relative abundance of *Escherichia Coli* among CEHC, CECRA and CECRC in CE subjects (*P*_CHEC-CECRA_=0.015, *P*_CEHC-CECRC_=0.0048). (d) The difference of the relative abundance of pks^+^ *Escherichia Coli* among Normal, CL and CE patients (*P*_Normal-CL_ < 0.0001, *P*_Normal-CE_ < 0.0001). (e) The difference of the relative abundance of pks^+^ *Escherichia Coli* among HC, CRA and CRC patients(*P*_HC-CRA_ < 0.0001,*P*_HC-CRC_ < 0.0001). (f) The difference of the relative abundance of pks^+^ *Escherichia Coli* among CEHC, CECRA and CECRC in CE subjects (*P*_CEHC-CECRA_ < 0.0001, *P*_CEHC-CECRC_< 0.0001). * *P*<0.05, ** *P*<0.01, *** *P*<0.001,**** *P* < 0.0001 *CEHC: patients with cholecystectomy and a normal colonoscope result; CECRA: patients with cholecystectomy and colorectal adenoma; CECRC: patients with cholecystectomy and colorectal cancer;

**Figure 4.**
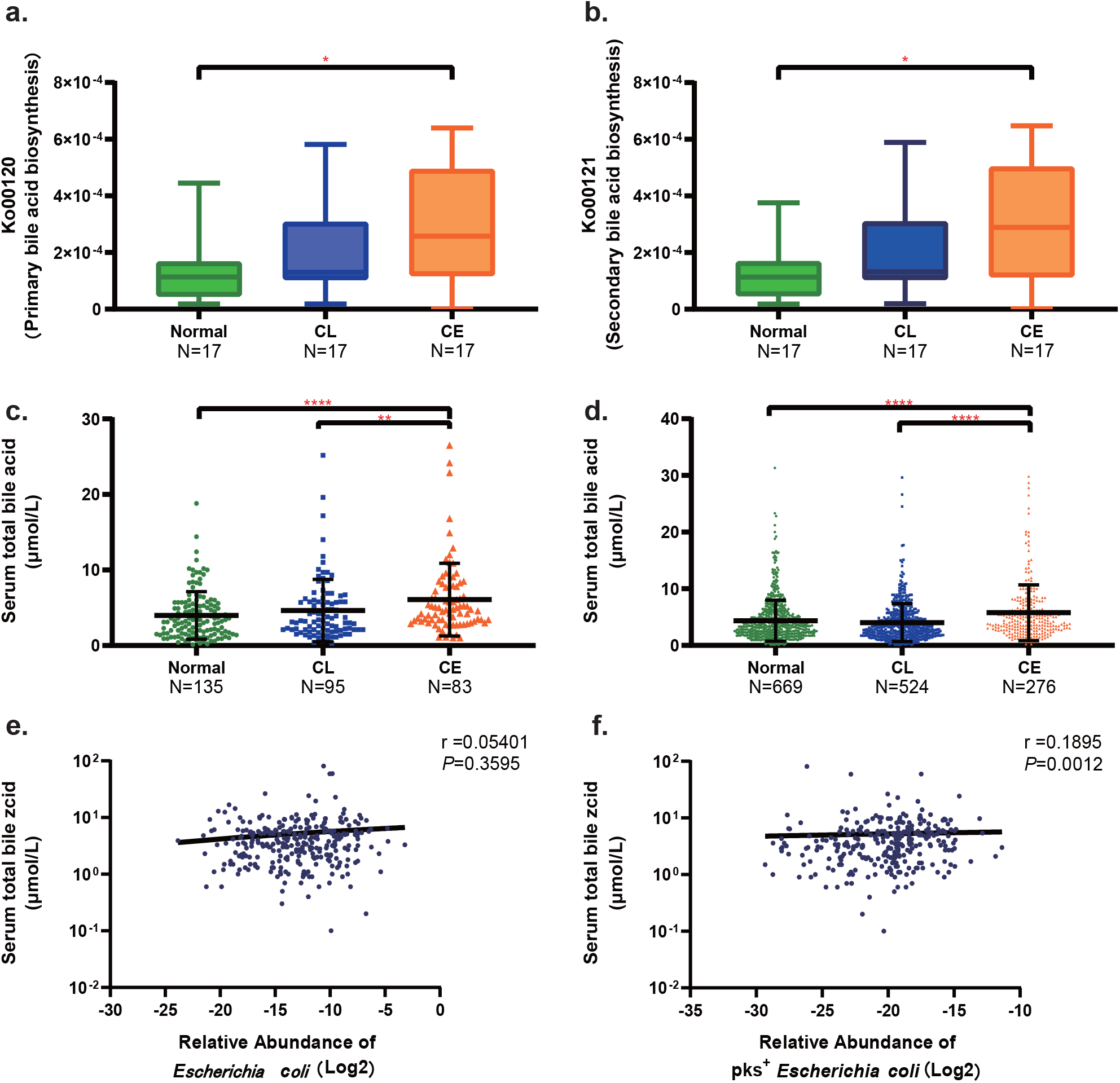
The difference of bile acid metabolism among Normal, CL and CE patients. (a)-(b) The relative abundance of Ko00120 (primary bile acid biosynthesis, a) and Ko00121 (secondary bile acid biosynthesis, b) through KEGG function prediction analysis by Tax4Fun in discovery cohort (Ko00120: *P*_Normal-CE_=0.024, Ko00121: *P*_Normal-CE_=0.027). (c)-(d) The level of serum total bile acid in validation cohort(c) (*P*_Normal-CE_<0.0001, *P*_CL-CE_=0.0035) and in database analysis(d) (*P*_Normal-CE_<0.0001, *P*_CL-CE_<0.0001). (e) Spearman correlation between the level of serum total bile acid and the relative abundance of *Escherichia Coli* (r=0.054, *P*=0.36). (f) Spearman correlation between the level of serum total bile acid and the relative abundance of pks^+^ *Escherichia Coli* (r=0.1895, *P*=0.0012). * *P*<0.05, ** *P*<0.01, *** *P*<0.001,**** *P* < 0.0001

All the other statistical analyses were performed using IBM SPSS STATISTICS (V.26.0) and GraphPad Prism (V.8.0). For continuous variables, Wilcoxon signed-rank test, Mann–Whitney U-test and t-test were used, and datas were expressed as mean ± standard deviation (SD). For categorical variables, the *χ*2 test was used and the datas were described as percentile. Spearman’s rank test and linear regression were used for correlation test. For all these comparisons, P<0.05 was considered statistically significant.

All the data, analytic methods, and study materials will be available to other researchers upon publication and upon request for peer review. The data that support the findings of this study are available in https://www.ncbi.nlm.nih.gov/sra/PRJNA895730, reference number PRJNA895730, and within the article and its supplementary materials.

## Results

### Patients baseline characteristics and quality control

To explore the differences of gut microbiota among patients with normal gallbladder (Normal), patients with cholelithiasis (CL) or with cholecystectomy (CE), we conducted a retrospective study recruited 463 volunteers in Renji Hospital. The discovery cohort was established to explore the differences of gut microbiota through 16S rRNA Illumina MiSeq sequencing. The validation cohort, which aimed to verify the results of sequencing in a larger sample size, included all 463 volunteers. There was no significant statistical difference in baseline categorical data including gender, age and body mass index (BMI) among Normal, CL and CE patients both in discovery and validation cohorts (Supplementary Table 2). Besides, no difference of tumor characteristics was found among patients in validation cohort with different gallbladder conditions (Supplementary Table 3).

To further explore the differences of tumor characteristics and bile acid metabolism among Normals, CLs and CEs, we also established a database analysis including 1,481 CRA and CRC patients (675 Normals, 526 CLs and 280 CEs) with different gallbladder conditions. There was also no significant statistical difference in baseline categorical data (Supplementary Table 2). The proportion of the CL and CE patients was 35.52% and 18.91%, which was significantly higher than population incidence(1). Among CRA patients, the occurrence of dysplasia in CE group was higher than that in the other groups (Normal vs CL vs CE: 21.39% vs 27.02% vs 36.14%, P=0.014, Supplementary Table 4). Among those patients with CRC, CL or CE patients were more likely to develop a larger adenocarcinoma (P<0.001) or a tumor with a later stage (P=0.002) (Supplementary Table 4).

After screening, 2,325,881 sequences belonged to 712 OTUs were acquired through the 16S rRNA sequencing of 51 stool specimens in the discovery cohort.

### The significant enrichment of *Escherichia coli* in patients with cholelithiasis or cholecystectomy

In the discovery cohort, we compared the composition of stool microbiota among Normal, CL and CE patients through 16S rRNA sequencing. For α-diversity, both the richness and the diversity of CE feces microbiota is lower than Normal group according to the Ace (*P*_Normal-CE_=0.046), Shannon (*P*_Normal-CE_=0.039), Sobs (*P*_Normal-CE_=0.033,) and Chao (*P*_Normal-CE_=0.039) indexes (Supplementary Figure 3).

Compared with Normal group, we found a significant elevation of the abundance of *Escherichia-Shigella* (in genus level, *P*_Normal-CL_=0.013, *P*_Normal-CE_=0.042; Figure 2a), and *Escherichia coli* (in species level, *P*_Normal-CL_=0.013, *P*_Normal-CE_=0.042; Figure 2b) in CL and CE groups. LEfSe analysis also suggested that *Escherichia coli* was the dominant species of the feces microbiota of CL and CE patients (LDA_Normal-CL_=3.97, *P*_Normal-CL_=0.013; LDA_Normal-CL_=4.52, *P*_Normal-CE_=0.040; Figure 2c).

To verify the result of 16S rRNA sequencing, the relative abundance of *Escherichia coli* has been assessed via qPCR in validation cohort. We found that the abundance of *Escherichia coli* increased 3.43 folds in CL group (*P*<0.0001) and 3.29 folds in CE group (*P*<0.0001) compared with Normal group (Figure 3a), which was consistent with the sequencing results.

### The increasement of *Escherichia coli* in CRA and CRC patients

To explore the abundance difference of *Escherichia coli* among HC, CRA and CRC patients, we reallocated the 51 volunteers in discovery cohort according to the results of colonoscopy, surgery and histopathology. The enrichment of *Enterobacterales* (in order level, *P*_HC-CRA_=0.035, *P*_HC-CRC_=0.011; Supplementary Figure 4a), *Enterobacteriaceae* (in family level, *P*_HC-CRA_=0.035, *P*_HC-CRC_=0.011; Supplementary Figure 4b), *Escherichia-Shigella* (in genus level, *P*_HC-CRA_=0.045, *P* _HC-CRC_=0.0016; Figure 2d), and *Escherichia coli* (in species level, *P*_HC-CRA_=0.045, *P* _HC-CRC_=0.0016; Figure 2e) was significant in CRA and CRC group compared with health control (HC). The results of LEfSe analysis reached the same findings (LDA_HC-CRA_=4.61, *P*_HC-CRA_=0.043; LDA _HC-CRC_=4.12, *P* _HC-CRC_=0.0015; Figure 2f). According to those results we figured out that CRA and CRC patients had a higher abundance of *Escherichia coli* than HC volunteers.

In the validation cohort, the relative abundance of *Escherichia coli* in stool specimens of CRC patients was proved to be 2.73 folds higher than HC patients via qPCR (*P*=0.0007), while the relative abundance of *Escherichia coli* of CRA patients was 2.26 folds (*P*=0.0063, Figure 3b). Among CE patients, we found that relative abundance of *Escherichia coli* in CECRC was 5.83 folds higher than CEHC subgroup (*P*=0.0048; Figure 3c). While such difference was smaller (3.71 folds) in Normal subjects (*P*_NHC-NCRC_=0.0044; Supplementary Figure 5a). However, there was no significant difference of *Escherichia coli* in CL group (Supplementary Figure 5c). We thus postulate that the elevated abundance of *Escherichia coli* may increase the incidence of colorectal tumor in patients after cholecystectomy.

### The enrichment of pks^+^ *Escherichia coli* in patients with cholelithiasis or cholecystectomy

For further exploration, we found that pks^+^ *Escherichia coli*, which proved to be a pernicious microbiota contributing to CRC(20, 21), was enriched in the stool specimens of CE patients. In validation cohort, the relative abundance of pks^+^ *Escherichia coli* in feces specimens of CE patients was 6.06 folds higher than Normal people (*P*<0.0001; Figure 3d), while there was no significant difference of cdt^+^ *Escherichia coli* (Supplementary Figure 6a). Furthermore, among CE patients, relative abundance of pks^+^ *Escherichia coli* in CECRC was 69.07 folds higher than in CEHC subgroup (*P*<0.0001, Figure 3f). While such difference in Normal people was lower (7.73 folds, *P*_NHC-NCRC_=0.027; Supplementary Figure 5b). For CL patients, although the relative abundance of pks^+^ *Escherichia coli* in feces specimens was higher than Normal subjects (P<0.0001, Figure 3d), there was no significant difference of pks^+^ *Escherichia coli* among CLHC, CLCRA and CLCRC (Supplementary Figure 5d). Therefore, we infer that pks^+^ *Escherichia coli* is correlated to the higher incidence of colorectal adenoma and adenocarcinoma in CE people.

### The effect of bile acid and pks^+^ *Escherichia coli* on tumorigenesis in patients with cholecystectomy

To explore the possible mechanism, KEGG functional predictive analysis was applied in discovery cohort through Tax4Fun. The expression of pathway for the biosynthesis of both primary and secondary bile acid in CE patients is higher than Normal group (Ko00120 primary bile acid biosynthesis *P*=0.014; Ko00121 secondary bile acid biosynthesis *P*=0.010; Figure 4a–b).

According to the laboratory metrics of validation cohort, the significant higher level of serum total bile acid was seen in CE patients than in Normal patients (Normal vs CE: 3.25 vs 4.75, *P*<0.0001; Figure 4c). We then analyzed the level of serum total bile acid among those 1,481 patients in database, and also found the increase of total bile acid in CE patients (Normal vs CE: 3.40 vs 4.20, P<0.0001; Figure 4d). Among those 1,481 patients, 289 patients (203 Normals, 56 CLs and 30 CEs) had detail information about bile acid metabolism profile. We found that the amount of secondary bile acid, including deoxycholic acid (Normal vs CE: 186.4 vs 339.2, P=0.016) and lithocholic acid (Normal vs CE: 12.0 vs 13.9, P=0.023), and their combining forms were higher in CE patients than in Normal people (Glycodeoxycholic acid, Normal vs CE: 212.6 vs 580.4, P=0.0063; Glycolithocholic acid, Normal vs CE: 8.50 vs 16.19, P=0.0080; Taurodeoxycholic acid, Normal vs CE: 45.75 vs 117.1, P=0.0014; Taurocholic acid, Normal vs CE: 2.000 vs 2.375, P=0.019) (Supplementary Figure 8). We suggest that bile acid, especially secondary bile acid is associated with the higher incidence of colorectal tumors in CE patients.

Furthermore, we found a significant correlation between the relative abundance of pks^+^ *Escherichia coli* in feces specimens and the level of serum total bile acid (r=0.1895, 95%CI 0.07214-0.3016, *P*=0.0012, Figure 4f) in validation cohort. Therefore, the pks^+^ *Escherichia coli* was found related to the bile acid metabolism in digestive tract. We can thus postulate that both the changes of microbiota and bile acid were associated with the occurrence and development of CRA and CRC in CE patients.

## Discussion

Gallbladder functions, gut microbiota and tumorigenesis of the digestive tract were found to be closely related with one another(11, 13, 14). However, to the furthest of our knowledge, the gut microbiota characteristic of CL or CE patients and the relationship among gallbladder conditions, gut microbiota and CRC has not been systematically studied and completely revealed. In current study, we have added to the present knowledge that the characteristics of gut microbiota are different among people with different gallbladder conditions. *Escherichia coli*, especially pks^+^ *Escherichia coli*, was found to be correlated with the occurrence and development of colorectal tumors in CE patients. Furthermore, we figured out that the interaction of pks^+^ *Escherichia coli* and bile acid metabolism was associated with colorectal tumors in CE patients.

In discovery cohort, lower α-diversity of gut microbiota was found in CE patients, and such changes were proved to promote the malignant changes of colorectal epithelial cells(22, 23). Through 16S rRNA sequencing and qPCR, we found significant enrichment of *Escherichia coli* in the stool specimens of CL and CE patients. Those results were consistent with the study by Wu *et al* about gut microbiota of patients with gallstones, which figured out the increasement of *proteobacteria* in CL patients(24). However, the aforementioned study generally paid attention to CL people instead of CE patients, and no specific bacteria species was identified. The results of current study therefore provided a better understanding to such findings. The differences of gut microbiota between CL patients and Normal subjects, as well as between CL patients before and after cholecystectomy, had also been partly revealed by Nirit Keren(9). For our current study, the differences of gut microbiota were compared among three groups, including Normal, CL and CE patients through bacterial species level, which adds to the present knowledge.

According to previous investigations, *Escherichia coli* was found more likely to be colonized in intestinal tissues and stool specimens of CRC patients(16), which is consistent with the results of current study. Some specific types of *Escherichia coli*, such as pks^+^ *Escherichia coli*(20, 21) or cdt^+^ *Escherichia coli*(25) were proved to contribute to CRC. It has been reported that pks^+^ *Escherichia coli* induces chromosomal instability and DNA damage through colibactin, which promotes epithelial cell proliferation and tumor invasion(26). Cdt^+^ *Escherichia coli* can produce cytolethal distending toxin, cause damage to host DNA and promote the occurrence and progression of CRC(25). In current study, the higher relative abundance of pks^+^ *Escherichia coli* in CL and CE patients were detected through qPCR in validation cohort. In CE patients, the relative abundance of pks^+^ *Escherichia coli* in CECRC was 69.07 folds higher than CEHC. The results above suggested that the colonization and enrichment of pks^+^ *Escherichia coli* in intestinal tract was associated with the higher occurrence of CRA and CRC in CE patients. Nevertheless, despite that higher abundance of pks^+^ *Escherichia coli* was found within CL patients, we did not find significant difference among HC, CRA and CRC subgroups.

It has been recognized that gut microbiota plays a crucial part in bile acid metabolism(27, 28). Meanwhile, the composition of intestinal bile acid can also change the structure of gut microbiota(9, 29). However, few studies have focused on the synergetic effect of gut microbiota and bile acid during the occurrence and development of CRA and CRC. In order to explore the possible mechanism, KEGG functional predictive analysis was utilized in our current study through Tax4Fun according to Silva database. The significant higher expression of Ko00120 and Ko00121, which was related to the biosynthesis of primary and secondary bile acids, was found in CE patients. In both validation cohort and database analysis, we found that the CE patients had a higher level of serum total bile acids than Normal people. Additionally, lithocholic acid and deoxycholic acid, which are the main component of secondary bile acid, together with their combining forms were found increased in CE patients. Previous studies had figured out the changes of the composition of intestinal bile acids and the enrichment of secondary bile acids in CE patients(11, 12), which were consistent to our findings. The tumorigenic activity of secondary bile acids has been reported in large number of experimental studies(11). Lithocholic acid and deoxycholic acid were proved to stimulate the proliferation of intestinal epithelial cells, promote DNA fragmentation and oxidative damage and would induce or promote the colorectal tumor development(11). Hence, we conjectured that the bile acids, especially secondary bile acids play important roles in the occurrence and development of CRC in CE patients. Furthermore, we found that the relative abundance of pks^+^ *Escherichia coli* in feces specimens was positively correlated with serum total bile acid level through Spearman’s rank correlation coefficient (*r*= 0.1895, *P*=0.0012). We thus postulate that the combination of pks^+^ *Escherichia coli* and bile acid are related to the higher incidence of colorectal adenoma and adenocarcinoma in people after cholecystectomy.

Moreover, we also figured out that CE patients would have more chance to get CRA with dysplasia or CRC with a larger size or later stage. Hence, surgical indications should be more carefully evaluated before cholecystectomy. For patients who have other high-risk factors of CRC, such as family history, smoking or history of CRA(30), risk of postoperative colorectal cancer shall be put into consideration. For those patients who are planned for cholecystectomy, surgeons shall make informed consent about their higher risk of CRA and CRC after surgery and suggest regular colonoscopy examination or FIT test after surgery. According to our findings, the regulation of gut microbiota and bile acid may be a new target for the prevention and treatment of CRA and CRC patients after cholecystectomy. Changing the composition and metabolism of gut microbiota and bile acid through changing diet habits or medications, such as probiotics, antibiotics or other ways that can regulate the intestinal bile acid pool, may reduce the incidence of CRA and CRC in CE patients. Besides, the enrichment of *Escherichia coli* and pks^+^ *Escherichia coli* were also found in CL patients, and pks^+^ *Escherichia coli* was reported to stimulate the malignant change of intestinal epithelium. Although no significant difference was found among CLHC, CLCRA and CLCRC patients, we also recommend earlier and more frequent colonoscopies or FIT tests among CL patients.

### Limitations

The study has several limitations. First, all volunteers were recruited in one single tertiary center. Multi-center studies are required to confirm the aforementioned conclusions among people in different countries and ethnicities, with different life backgrounds and experiences. Secondly, prospective studies should be executed to track the change process of gut microbiota and bile acid in people after cholecystectomy. Further researches are necessary to explore and validate the mechanism and causality between pks^+^ *Escherichia coli* and bile acid in the development of colorectal cancer among CE patients.

### Conclusions

Our study revealed the differences of gut microbiota among people with normal gallbladder and patients with cholelithiasis or cholecystectomy. We figured out that the relative abundance of *Escherichia coli*, especially pks^+^ *Escherichia coli*, increased in CL and CE patients. We also found that the interaction of pks^+^ *Escherichia coli* and bile acids was associated with the elevation of occurrence of colorectal adenoma and adenocarcinoma in people after cholecystectomy. The findings of current study provided a novel insight into the pathogenesis of colorectal tumor in patients have gallbladder conditions and hinted a new pathway for future treatment and prevention of colorectal tumor in patients with cholelithiasis or cholecystectomy.

## Abbreviations

BMI: body mass index
CE: patients with cholecystectomy
CEHC: patients with cholecystectomy and a normal colonoscope result
CECRA: patients with cholecystectomy and colorectal adenoma
CECRC: patients with cholecystectomy and colorectal cancer
CL: patients with cholelithiasis
CLHC: patients with cholelithiasis and a normal colonoscope result
CLCRA: patients with cholelithiasis and colorectal adenoma
CLCRC: patients with cholelithiasis and colorectal cancer
CRA: colorectal adenoma
CRC: colorectal cancer
Ct: cycle threshold
CT: computed tomography
FIT: fecal immunochemical tests
HC: people with normal result of colonoscopy
KEGG: Kyoto Encylopaedia of Genes and Genomes
LDA: linear discriminant analysis
MRI: magnetic resonance imaging
NCRA: patients with normal gallbladder and colorectal adenoma
NCRC: patients with normal gallbladder and colorectal cancer
NHC: volunteers both with a normal gallbladder and a normal colonoscope result), Normal(patients with normal gallbladder
OTU: operational taxonomic unit
PCR: polymerase chain reaction
qPCR: quantitative real-time polymerase chain reaction).

## Author Contributors

SY Pan and JY Fang designed this study and were responsible for data analysis and interpretation. SY Pan also took part in subject recruitment, sample collection, experiment and manuscript writing. CB Zhou assisted on data analysis, interpretation and manuscript revision. JY Fang also provided financial support for this study. JW Deng, YL Zhou and ZH Liu were responsible for subject recruitment and sample collection. All authors reviewed the draft of this manuscript and approved the final version.

## Declaration of interests

The authors declared no competing interests.

## Acknowledgements

This study was supported by National Key R&D Program of China (2020YFA0509200) (2016YFC0906002) and grants from the National Natural Science Foundation of China (81830081, 81972203, 31970718, 82070570, 82203224). The project was also partially supported by Shanghai Municipal Health Commission, Collaborative Innovation Cluster Project (2019CXJQ02) and Shanghai Sailing Program (21YF1425600), China Postdoctoral Science Foundation (2022M712124). We are grateful to all members of the group at the Shanghai Institute of Digestive Disease for providing assistance in sample collection and data analysis. We thank all the patients and their families for participating in this study.

## Patient consent

Obtained.

## Ethical approval

This study was approved by Ethics Committee of Shanghai Jiao-Tong University School of Medicine, Renji Hospital (Shanghai, China)(#2022-065-B).

## References

1. European Association for the Study of the Liver. Electronic address eee. 2016. EASL Clinical Practice Guidelines on the prevention, diagnosis and treatment of gallstones. J Hepatol 65:146–181.

2. Tazuma S, Unno M, Igarashi Y, Inui K, Uchiyama K, Kai M, Tsuyuguchi T, Maguchi H, Mori T, Yamaguchi K, Ryozawa S, Nimura Y, Fujita N, Kubota K, Shoda J, Tabata M, Mine T, Sugano K, Watanabe M, Shimosegawa T. 2017. Evidence-based clinical practice guidelines for cholelithiasis 2016. J Gastroenterol 52:276–300.

3. Gutt C, Schlafer S, Lammert F. 2020. The Treatment of Gallstone Disease. Dtsch Arztebl Int 117:148–158.

4. Chiong C, Cox MR, Eslick GD. 2012. Gallstones are associated with colonic adenoma: a meta-analysis. World J Surg 36:2202–9.

5. Zhang Y, Liu H, Li L, Ai M, Gong Z, He Y, Dong Y, Xu S, Wang J, Jin B, Liu J, Teng Z. 2017. Cholecystectomy can increase the risk of colorectal cancer: A meta-analysis of 10 cohort studies. PLoS One 12:e0181852.

6. Siddiqui AA, Kedika R, Mahgoub A, Patel M, Cipher DJ, Bapat V. 2009. A previous cholecystectomy increases the risk of developing advanced adenomas of the colon. South Med J 102:1111–5.

7. Chen YK, Yeh JH, Lin CL, Peng CL, Sung FC, Hwang IM, Kao CH. 2014. Cancer risk in patients with cholelithiasis and after cholecystectomy: a nationwide cohort study. J Gastroenterol 49:923–31.

8. Ward HA, Murphy N, Weiderpass E, Leitzmann MF, Aglago E, Gunter MJ, Freisling H, Jenab M, Boutron-Ruault MC, Severi G, Carbonnel F, Kuhn T, Kaaks R, Boeing H, Tjonneland A, Olsen A, Overvad K, Merino S, Zamora-Ros R, Rodriguez-Barranco M, Dorronsoro M, Chirlaque MD, Barricarte A, Perez-Cornago A, Trichopoulou A, Bamia C, Lagiou P, Masala G, Grioni S, Tumino R, Sacerdote C, Mattiello A, Bueno-de-Mesquita B, Vermeulen R, Van Gils C, Nystrom H, Rutegard M, Aune D, Riboli E, Cross AJ. 2019. Gallstones and incident colorectal cancer in a large pan-European cohort study. Int J Cancer 145:1510–1516.

9. Keren N, Konikoff FM, Paitan Y, Gabay G, Reshef L, Naftali T, Gophna U. 2015. Interactions between the intestinal microbiota and bile acids in gallstones patients. Environ Microbiol Rep 7:874–80.

10. Thomas LA, Veysey MJ, Bathgate T, King A, French G, Smeeton NC, Murphy GM, Dowling RH. 2000. Mechanism for the transit-induced increase in colonic deoxycholic acid formation in cholesterol cholelithiasis. Gastroenterology 119:806–15.

11. Cao H, Xu M, Dong W, Deng B, Wang S, Zhang Y, Wang S, Luo S, Wang W, Qi Y, Gao J, Cao X, Yan F, Wang B. 2017. Secondary bile acid-induced dysbiosis promotes intestinal carcinogenesis. Int J Cancer 140:2545–2556.

12. Zhang F, Duan Y, Xi L, Wei M, Shi A, Zhou Y, Wei Y, Wu X. 2018. The influences of cholecystectomy on the circadian rhythms of bile acids as well as the enterohepatic transporters and enzymes systems in mice. Chronobiol Int 35:673–690.

13. Wirbel J, Pyl PT, Kartal E, Zych K, Kashani A, Milanese A, Fleck JS, Voigt AY, Palleja A, Ponnudurai R, Sunagawa S, Coelho LP, Schrotz-King P, Vogtmann E, Habermann N, Nimeus E, Thomas AM, Manghi P, Gandini S, Serrano D, Mizutani S, Shiroma H, Shiba S, Shibata T, Yachida S, Yamada T, Waldron L, Naccarati A, Segata N, Sinha R, Ulrich CM, Brenner H, Arumugam M, Bork P, Zeller G. 2019. Meta-analysis of fecal metagenomes reveals global microbial signatures that are specific for colorectal cancer. Nat Med 25:679–689.

14. Wong SH, Yu J. 2019. Gut microbiota in colorectal cancer: mechanisms of action and clinical applications. Nat Rev Gastroenterol Hepatol 16:690–704.

15. Rubinstein MR, Wang X, Liu W, Hao Y, Cai G, Han YW. 2013. Fusobacterium nucleatum promotes colorectal carcinogenesis by modulating E-cadherin/beta-catenin signaling via its FadA adhesin. Cell Host Microbe 14:195–206.

16. Bonnet M, Buc E, Sauvanet P, Darcha C, Dubois D, Pereira B, Dechelotte P, Bonnet R, Pezet D, Darfeuille-Michaud A. 2014. Colonization of the human gut by E. coli and colorectal cancer risk. Clin Cancer Res 20:859–67.

17. Fais T, Delmas J, Barnich N, Bonnet R, Dalmasso G. 2018. Colibactin: More Than a New Bacterial Toxin. Toxins (Basel) 10.

18. Remacle AG, Shiryaev SA, Strongin AY. 2014. Distinct interactions with cellular E-cadherin of the two virulent metalloproteinases encoded by a Bacteroides fragilis pathogenicity island. PLoS One 9:e113896.

19. Amin MB, Greene FL, Edge SB, Compton CC, Gershenwald JE, Brookland RK, Meyer L, Gress DM, Byrd DR, Winchester DP. 2017. AJCC Cancer Staging Manual. 8th ed. New York,. NY: Springer doi:10.3322/caac. 21388:252–254.

20. Wilson MR, Jiang Y, Villalta PW, Stornetta A, Boudreau PD, Carra A, Brennan CA, Chun E, Ngo L, Samson LD, Engelward BP, Garrett WS, Balbo S, Balskus EP. 2019. The human gut bacterial genotoxin colibactin alkylates DNA. Science 363.

21. Nougayrede JP, Homburg S, Taieb F, Boury M, Brzuszkiewicz E, Gottschalk G, Buchrieser C, Hacker J, Dobrindt U, Oswald E. 2006. Escherichia coli induces DNA double-strand breaks in eukaryotic cells. Science 313:848–51.

22. Kriss M, Hazleton KZ, Nusbacher NM, Martin CG, Lozupone CA. 2018. Low diversity gut microbiota dysbiosis: drivers, functional implications and recovery. Curr Opin Microbiol 44:34–40.

23. Ahn J, Sinha R, Pei Z, Dominianni C, Wu J, Shi J, Goedert JJ, Hayes RB, Yang L. 2013. Human gut microbiome and risk for colorectal cancer. J Natl Cancer Inst 105:1907–11.

24. Wu T, Zhang Z, Liu B, Hou D, Liang Y, Zhang J, Shi P. 2013. Gut microbiota dysbiosis and bacterial community assembly associated with cholesterol gallstones in large-scale study. BMC Genomics 14:669.

25. Grasso F, Frisan T. 2015. Bacterial Genotoxins: Merging the DNA Damage Response into Infection Biology. Biomolecules 5:1762–82.

26. Pleguezuelos-Manzano C, Puschhof J, Rosendahl Huber A, van Hoeck A, Wood HM, Nomburg J, Gurjao C, Manders F, Dalmasso G, Stege PB, Paganelli FL, Geurts MH, Beumer J, Mizutani T, Miao Y, van der Linden R, van der Elst S, Genomics England Research C, Garcia KC, Top J, Willems RJL, Giannakis M, Bonnet R, Quirke P, Meyerson M, Cuppen E, van Boxtel R, Clevers H. 2020. Mutational signature in colorectal cancer caused by genotoxic pks(+) E. coli. Nature 580:269–273.

27. Urdaneta V, Casadesus J. 2017. Interactions between Bacteria and Bile Salts in the Gastrointestinal and Hepatobiliary Tracts. Front Med (Lausanne) 4:163.

28. Molinero N, Ruiz L, Sanchez B, Margolles A, Delgado S. 2019. Intestinal Bacteria Interplay With Bile and Cholesterol Metabolism: Implications on Host Physiology. Front Physiol 10:185.

29. Sauter GH, Moussavian AC, Meyer G, Steitz HO, Parhofer KG, Jungst D. 2002. Bowel habits and bile acid malabsorption in the months after cholecystectomy. Am J Gastroenterol 97:1732–5.

30. Tao S, Hoffmeister M, Brenner H. 2014. Development and validation of a scoring system to identify individuals at high risk for advanced colorectal neoplasms who should undergo colonoscopy screening. Clin Gastroenterol Hepatol 12:478–85.

